# Gadolinium-based nanoparticles AGuIX and their combination with ionizing radiation trigger AMPK-dependent proinflammatory reprogramming of tumor-associated macrophages

**DOI:** 10.1101/2024.01.12.575217

**Authors:** Zeinaf Muradova, Désirée Tannous, Ali Mostefa-Kara, Thanh Trang Cao-Pham, Constance Lamy, Sophie Broutin, Angelo Paci, Sandrine Dufort, Tristan Doussineau, François Lux, Olivier Tillement, Géraldine Le Duc, Awatef Allouch, Jean-Luc Perfettini

## Abstract

**Background:** Tumor-associated macrophages (TAMs) are essential components of the inflammatory microenvironment of tumors and are associated with poor clinical outcomes in the majority of cancers. TAMs mainly exhibit anti-inflammatory functions that promote and support the tissue remodeling, the immune suppression and the tumor growth. Regarding their plasticity, the functional reprogramming of anti-inflammatory TAMs into proinflammatory phenotype recently emerged as a therapeutic opportunity to improve the effectiveness of anticancer treatments such as radiotherapy.

**Results:** Here we show that gadolinium-based nanoparticles AGuIX alone and in combination with ionizing radiation (IR) induce in a dose-dependent manner, the accumulation of DNA double strand breaks, an Ataxia telangiectasia mutated (ATM)-dependent DNA-damage response, an increased expression of the Interferon regulatory factor 5 (IRF5) and the release of proinflammatory cytokines from targeted macrophages, thus directing their proinflammatory reprogramming. This process is associated with the activating phosphorylation of the Adenosine Monophosphate (AMP) activated protein kinase on threonine 172 (AMPKT172*) and the fragmentation of mitochondria. Furthermore, we demonstrate that the inactivation of AMPK reduces the mitochondrial fragmentation and the proinflammatory reprogramming of macrophages detected in response to AGuIX and their combination with IR. These results reveal that the AMPK-dependent regulation of mitochondrial fragmentation plays a central role during the proinflammatory reprogramming of macrophages. Accordingly, a positive correlation between AMPKT172* and proinflammatory activation of TAMs is detected following IR+AGuIX combination in syngeneic mouse model of colorectal cancer.

**Conclusions:** Altogether, our results identify a novel signaling pathway elicited by AGuIX and their combined treatment with IR, that targets macrophage polarization, skews macrophage functions toward the proinflammatory phenotype and may enhance the effectiveness of radiotherapy.

## Background

The tumor microenvironment (TME) is an immunosuppressive niche supporting the cancer progression and the immune escape that recently emerged as promising therapeutic target for cancer treatment [1, 2]. Tumor-associated macrophages (TAMs) account for the most abundant myeloid cells in the TME [3] and support both cancer progression and immune evasion [4–6]. One main feature of TAMs is their high plasticity and ability to adapt and reprogram their biological functions to environmental signals [7]. TAMs demonstrate anti-inflammatory properties that stimulate angiogenesis, cancer cell invasion and metastatic dissemination. TAMs also contribute to immunosuppression in the TME by recruiting regulatory T cells and myeloid-derived suppressor cells (MDSCs), inhibiting dendritic cell maturation and/or expressing inhibitory innate and immune adaptive checkpoint proteins (such as SIRPα, PD1 or PDL1) [8, 9]. Histological detection of TAMs predicts treatment response and is associated with poor clinical outcomes in the majority of solid cancers and hematological malignancies [10, 11]. Targeting the functional reprogramming of TAMs has been proposed to improve the efficacy of anticancer treatments including radiotherapy [8, 12]. Highly plastic, anti-inflammatory TAMs can thus be functionally reprogrammed into proinflammatory phenotype to establish a tumoricidal microenvironment and to support the development of long-lasting specific anti-tumor immune response. Recently, we demonstrated that ionizing radiation (IR) reprograms pro-tumorigenic macrophages toward a proinflammatory phenotype. The IR-mediated macrophage reprogramming that we identified requires the activation of the NADPH oxidase 2 (NOX2), the production of reactive oxygen species (ROS), the activating phosphorylation of the Ataxia telangiectasia mutated (ATM) kinase on serine 1981 (ATMS1981*) and the transcriptional activity of the Interferon regulatory factor 5 (IRF5) [13]. Briefly, IR triggered DNA damage into the nuclei of targeted macrophages, leading to a DNA damage response that is directed by the ATM kinase. In response to IR, the ATM kinase, which is activated by phosphorylation on serine 1981 (ATMS1981*), controls the macrophage phenotypic conversion from anti-inflammatory to proinflammatory phenotype by regulating mRNA expression of IRF5. To further characterize this process, we deciphered upstream signaling pathways and detected that ROS are produced in response to IR. Interestingly, we revealed that NOX2, whose expression is also increased after IR, is involved in ROS production and directed the IR-mediated proinflammatory macrophage reprogramming. Moreover, the detection of this signaling pathway on patient biopsies predicts a good tumor response to preoperative radiotherapy in locally advanced rectal cancer [13]. Altogether, these results convinced us that the identification of novel therapeutic strategy that may enhance the ability of IR to stimulate this signaling pathway and to reprogram TAMs would improve radiotherapy efficacy.

Several applications of nanomedecine (such as radioisotope-labeled or metallic nanoparticles) have been developed to improve the therapeutic index of radiation therapy. Nanomaterials were initially used as drug delivery platforms or as contrast agents, to enhance the magnetic resonance imaging (MRI) contrast of tumors and to develop MRI-guided radiotherapy, but also as radiosensitizers, to improve the delivery and the deposition of radiation doses into tumor sites [14–18]. Considering the ability of metallic nanoparticles containing high-Z atoms to increase proportionally to their atomic number, the radiation dose absorbed by tissue, gadolinium-based nanoparticles AGuIX have been extensively investigated for their potential to improve radiotherapy. Under exposure to IR, AGuIX produce photons and Auger electrons that improve the nanoscale dose rate deposition in the vicinity of the nanoparticles into the tumors, enhance the DNA double-strand break damage and the production of ROS, and lead to the destruction of numerous tumors (e.g. melanoma, glioblastoma, breast and lung carcinomas)[19–21]. Nanomedicine has been recently proposed to amplify antitumor immune response and to sensitize tumors to RT and/or immunotherapies [22, 23]. Although the combination of AGuIX with IR (AGuIX+IR) stimulated a growing interest for cancer treatment (brain metastases: NCT02820454, NCT03818386 and NCT04899908; lung and pancreatic cancer: NCT04789486, cervix cancer: NCT03308604 and glioblastoma: NCT04881032), the immune response induced by AGuIX+IR combination is poorly understood. Here, we revealed *in vitro* that AGuIX alone and their combination with IR induced DNA double-strand break damage into the nuclei of treated macrophages and triggered their proinflammatory reprogramming in an adenosine monophosphate (AMP) activated protein kinase (AMPK)-dependent manner. Strikingly, the number of TAMs showing AMPK activation or undergoing proinflammatory activation in response to IR and AGuIX combination is significantly enhanced in a syngeneic mouse model of colorectal cancer. Altogether, these results demonstrated that AGuIX+IR combination could be used to convert anti-inflammatory TAMs into proinflammatory macrophages and to improve the efficacy of radiotherapy.

## Results

### Gadolinium-based nanoparticles AGuIX and their combination with ionizing radiation trigger DNA double-strand breaks and an ATM-dependent DNA damage response in human and murine macrophages

We initially demonstrated that DNA damage plays a central role during proinflammatory macrophage activation [13]. Considering the ability of the combination of AGuIX with IR to increase dose deposition and consequently enhance the accumulation of DNA damage in treated cancer cells [24], we hypothesized that AGuIX+IR combination may also induce DNA damage in TAMs and strengthen their ability to be converted into proinflammatory macrophages in response to IR. Using confocal microscopy, we first studied the induction of DNA double-strand breaks in phorbol-12-myristate 13-acetate (PMA)-differentiated human THP1 macrophages that were exposed to a single dose of 0.2 Gy in the presence of different concentrations of AGuIX. As expected, thirty minutes after IR, nuclear foci containing the phosphorylated form of the histone variant H2AX on serine 139 (H2AXS139*) (also known as γ-H2AX foci) were detected in treated macrophages (Figs. 1a-1c). Moreover, the combination of IR with 200 nM AGuIX significantly enhanced the frequency of macrophages showing the nuclear accumulation of γ-H2AX foci, as compared to control or irradiated macrophages (Figs. 1b and 1c). We noticed that these nuclear γ-H2AX foci persisted until 48 hours after the combined treatment of IR with 200 nM AGuIX (Fig. 1c). Interestingly, we observed that higher concentrations of AGuIX (0.6 mM and 1.2 mM) allowed after 1 hour of treatment, the nuclear accumulation of γ-H2AX foci in treated macrophages (Fig. 1f). The frequency of PMA-differentiated human THP1 macrophages showing nuclear γ-H2AX foci and the size of these nuclear foci increased when macrophages were irradiated with a single dose of 0.2 Gy in presence of 0.6 mM AGuIX or 1.2 mM AGuIX (Figs. 1d-1f). These results were confirmed on murine RAW264.7 macrophages exposed to a single dose of 0.2 Gy IR in presence of 0.6 mM AGuIX or 1.2 mM AGuIX (Fig. 1g), thus revealing the ability of AGuIX+IR combination to increase the dose deposition and the accumulation of DNA double-strand breaks in treated macrophages. We then analyzed the DNA damage response (DDR) elicited by these genomic alterations and determined whether the DDR is dependent of the Ataxia telangiectasia mutated (ATM) kinase, as we previously described [13]. We thus studied the activating phosphorylation of the kinase ATM on serine 1981 (ATMS1981*) in PMA-differentiated human THP1 macrophages (Fig. 1h) that were exposed to a single dose of 0.2 Gy IR in presence of different concentrations of AGuIX. Thirty minutes after treatments, we detected that 100 nM and 200 nM AGuIX induced ATMS1981* (Fig. 1h) and that the level of ATMS1981* phosphorylation is significantly increased after the treatment of PMA-differentiated human THP1 macrophages with AGuIX+IR combination, as compared to control cells (Fig. 1h). Murine RAW264.7 macrophages irradiated with a single dose of 0.2 Gy in presence of 0.6 mM AGuIX or 1.2 mM AGuIX also accumulated ATMS1981* in their nuclei and showed a significant increase in the frequency of macrophages showing nuclear ATMS1981* after 1 hour-treatment with AGuIX alone (0.6 mM AGuIX or 1.2 mM AGuIX) or with AGuIX+IR combination (Fig. 1i). Altogether, these results revealed the ability of AGuIX to direct DNA damage in macrophages and to enhance the amount of DNA damage and DDR in anti-inflammatory macrophages that were irradiated in the presence of AGuIX.

**Fig. 1.**
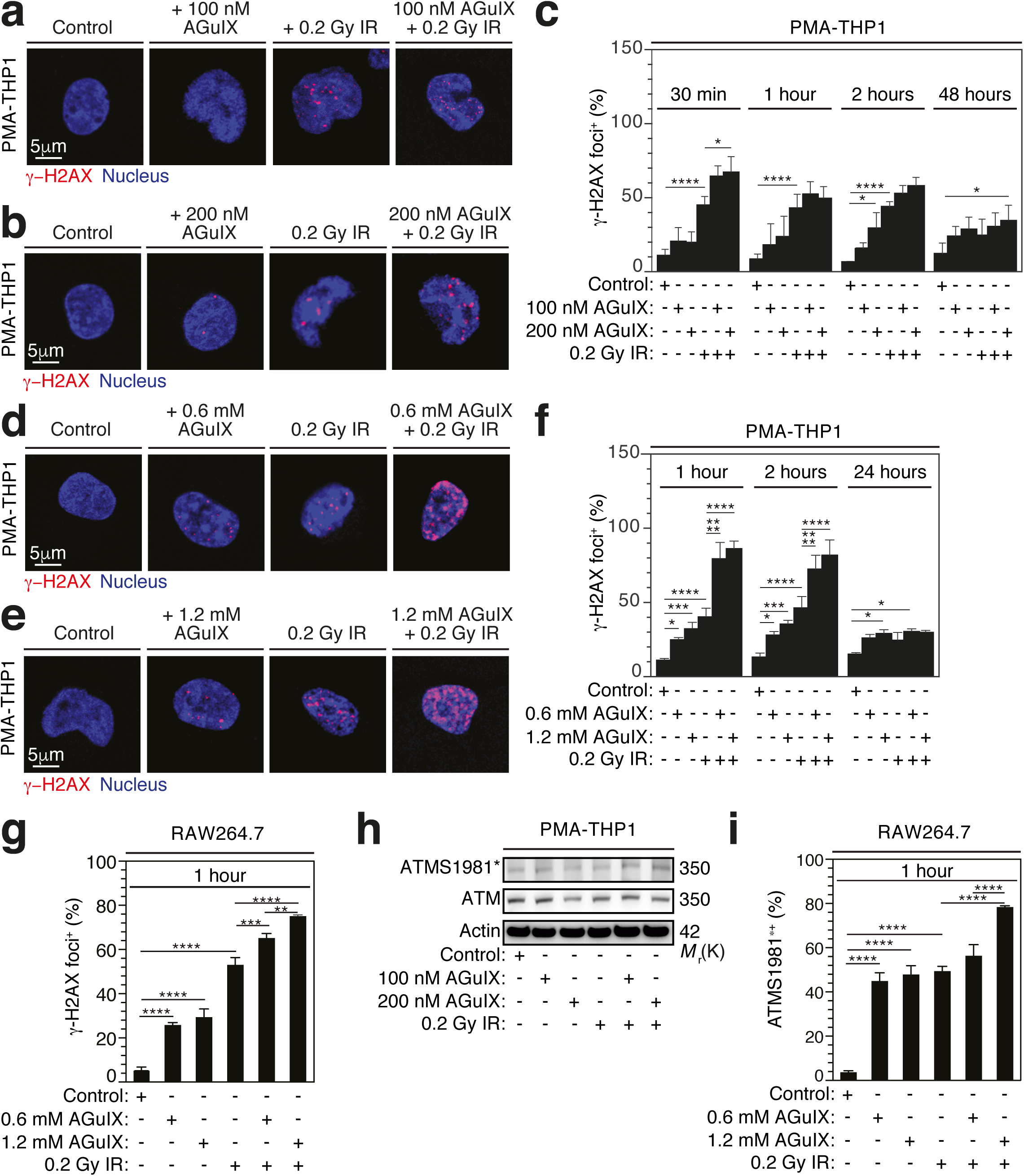
AGuIX alone and combined treatment with IR induces DNA damage and DDR in macrophages. **a**-**c** Confocal micrographs (**a**, **b**) and percentages (**c**) of PMA-differentiated human THP1 macrophages showing γ-H2AX foci after 30 minutes (**a**-**c**) and 1 hour, 2 hours or 48 hours (**c**) of treatment with control, 100 nM (**a**, **c**) or 200 nM (**b**, **c**) AGuIX, 0.2 Gy IR (**a-c**) and the combination of 0.2 Gy IR with 100 nM (**a**, **c**) or 200 nM (**b**, **c**) AGuIX. **d**-**f** Confocal micrographs (**d**, **e**) and percentages (**f**) of PMA-differentiated THP1 macrophages showing γ-H2AX foci after 1 hour (**d**-**f**) and 2 hours or 24 hours (**f**) of treatment with control, 0.6 mM (**d**, **f**) or 1.2 mM (**e**, **f**) AGuIX, 0.2 Gy IR (**d-f**) and the combination of 0.2 Gy IR with 0.6 mM (**d**, **f**) or 1.2 mM (**e**, **f**) AGuIX. **g** Percentages of murine RAW264.7 macrophages showing γ-H2AX foci at 1 hour after treatments with control, 0.6 mM AGuIX, 1.2 mM AGuIX, 0.2 Gy IR and the combination of 0.2 Gy IR with 0.6 mM or 1.2 mM AGuIX. **h** ATMS1981* and ATM expressions detected in PMA-differentiated THP1 macrophages after 30-minute treatment with control, 100 nM AGuIX, 200 nM AGuIX, 0.2 Gy IR and the combination of 0.2 Gy IR with 100 nM or 200 nM AGuIX. Actin is used as loading control. **i** Percentages of ATMS1981*^+^ cells detected on murine RAW264.7 macrophages treated as in (**g**). Data in **a**, **b**, **d**, **e** and **h** are representative of n=3 independent experiments. Data in **c**, **f**, **g** and **i** are means ± S.E.M from n=3 independent experiments. P-values (****P< 0.0001, ***P< 0.001, **P< 0.01 and *P<0.05) were calculated by using two-way ANOVA (**c**, **f**) or one-way ANOVA tests (**g**, **i**) with Tukey’s multiple comparisons.

### Gadolinium-based nanoparticles AGuIX and their combination with ionizing radiation favor the proinflammatory reprogramming of macrophages

Considering that we previously revealed that DDR and ATM dictate the proinflammatory macrophage phenotype [13], we then studied the impact of AGuIX and their combination with IR on macrophage functional reprogramming. Using fluorescent microscopy, we first determined the expression level of a proinflammatory marker of macrophage activation, the inducible nitric oxide synthase (iNOS) on PMA-differentiated human THP1 macrophages that were irradiated with 0.2 Gy in combination with different concentrations of AGuIX. Although PMA-treated human THP1 macrophages that were treated with 100 nM or 200 nM of AGuIX and/or irradiated with 0.2 Gy did not exhibit an increased expression of iNOS at 2 hours after irradiation, the frequency of iNOS^+^ macrophages significantly increased after 48 hours in presence of 100 nM or 200 nM AGuIX alone, 0.2 Gy irradiation or the combination of 200 nM AGuIX with 0.2 Gy irradiation (Figs. 2a-2c). Interestingly, we observed that PMA-differentiated human THP1 macrophages that were treated with 0.6 mM or 1.2 mM AGuIX also revealed a significant increase of iNOS expression levels, as compared to control macrophages (Figs. 2d-2f), thus revealing that the potential of AGuIX to trigger the proinflammatory phenotype of macrophages in absence of IR. As expected, the combination of 0.6 mM or 1.2 mM AGuIX with IR enhanced significantly the frequency of iNOS^+^ cells as compared to irradiated cells (Figs. 2d-2f). To confirm these results, we determined by western blot analysis the cellular expression of IRF5, a transcriptional factor of macrophage proinflammatory reprogramming, and the release into the supernatant of two cytokines, interleukins 1β (IL-1β) and 6 (IL-6) that we previously detected during IR-mediated proinflammatory macrophage reprogramming [13]. We observed that after 48 hours (Fig. 2g) or 24 hours (Fig. 2h) of irradiation, the combination of IR with 100 nM, 200 nM, 0.6 mM or 1.2 mM of AGuIX enhanced the expression of IRF5 in treated macrophages (Figs. 2g and 2h). These treatments were associated with the release of IL-1β, IL-6 in the supernatant of treated macrophages (IR + 100 nM AGuIX or IR + 200 nM AGuIX) (Fig. 2i). Interestingly, we detected that AGuIX alone (100 nM AGuIX or 200 nM AGuIX) increased the expression of IRF5 in PMA-treated human THP1 macrophages (Figs. 2g and 2h) and the secretion of proinflammatory cytokine IL-6 in absence of IR (Fig. 2i). Altogether, these results indicate that AGuIX prime anti-inflammatory macrophages for proinflammatory reprogramming and as a consequence, their combination with IR enhances the ability of irradiated macrophages to acquire a proinflammatory phenotype.

**Fig. 2.**
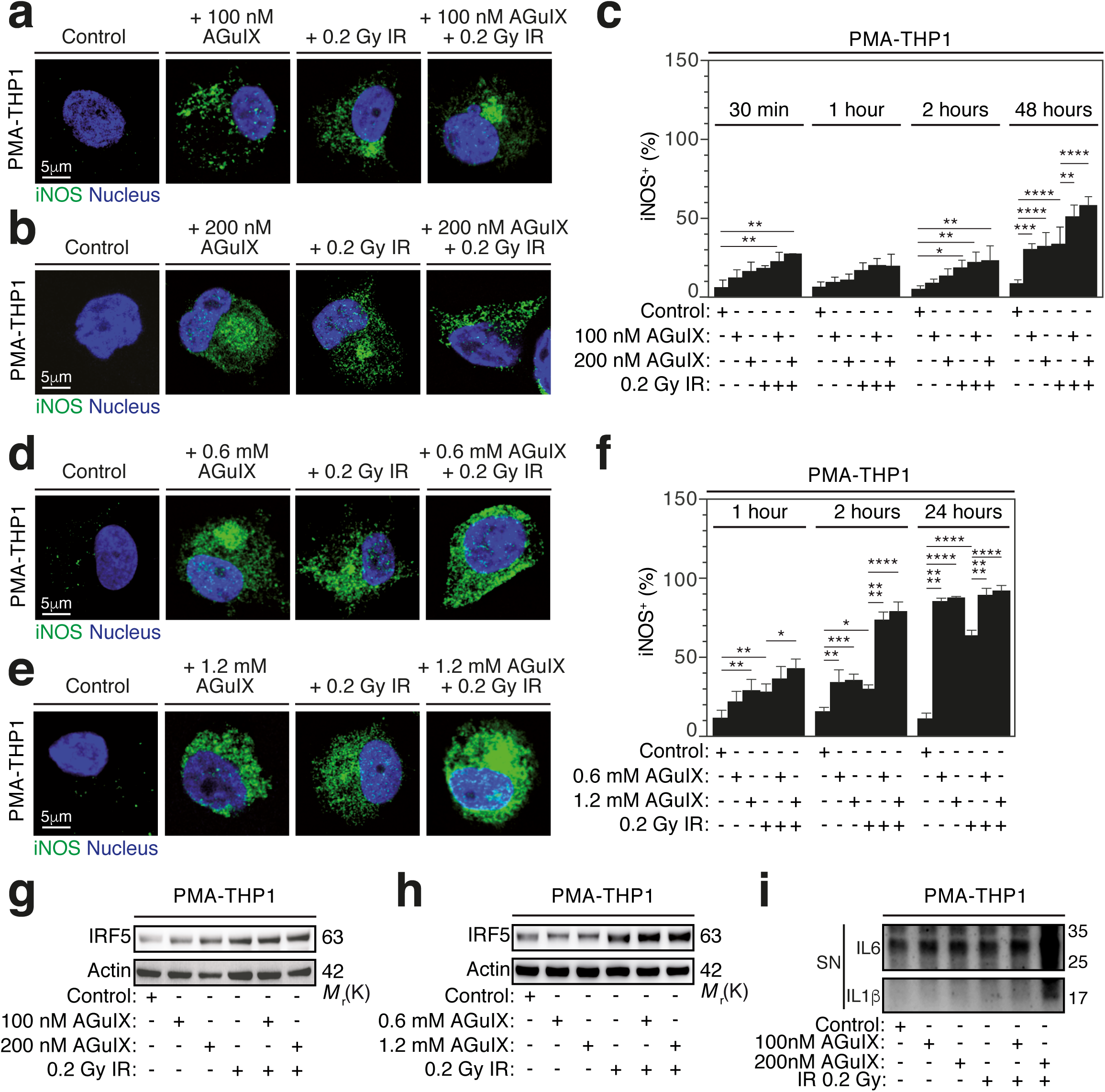
AGuIX alone and combined treatment with IR activates macrophages toward a proinflammatory phenotype. **a**-**c** Confocal micrographs (**a**, **b**) and percentages (**c**) of PMA-differentiated human THP1 macrophages expressing iNOS (iNOS^+^) detected after 30 minutes (**a**-**c**) and 1 hour, 2 hours or 48 hours (**c**) of treatment with control, 100 nM (**a**, **c**) or 200 nM (**b**, **c**) AGuIX, 0.2 Gy IR (**a-c**) and the combination of 0.2 Gy IR with 100 nM (**a**, **c**) or 200 nM (**b**, **c**) AGuIX. **d**-**f** Confocal micrographs (**d**, **e**) and percentages (**f**) of iNOS^+^ PMA-differentiated THP1 macrophages detected after 1 hour (**d**-**f**), 2 hours (**f**) or 24 hours (**f**) of treatment with control, 0.6 mM (**d**, **f**) or 1.2 mM (**e**, **f**) AGuIX, 0.2 Gy IR (**d-f**) and the combination of 0.2 Gy IR with 0.6 mM (**d**, **f**) or 1.2 mM (**e**, **f**) AGuIX. **g**-**i** IRF5 expression (**g**, **h**) and release of IL-1β and IL6 in the cell culture supernatant (SN) (**i**) detected after 48-hour treatment of PMA-differentiated THP1 macrophages with control, 100 nM AGuIX, 200 nM AGuIX, 0.2 Gy IR and the combination of 0.2 Gy IR with 100 nM or 200 nM AGuIX (**g**, **i**), or with control, 0.6 mM AGuIX, 1.2 mM AGuIX, 0.2 Gy IR and the combination of 0.2 Gy IR with 0.6 mM or 1.2 mM AGuIX (**h**). Actin is used as loading control in **g** and **h**. Data in **a**, **b**, **d**, **e**, **g**, **h** and **i** are representative of n=3 independent experiments are shown. Data in **c** and **f** are means ± S.E.M from n=3 independent experiments. P-values (****P< 0.0001, ***P< 0.001, **P< 0.01 and *P<0.05) were calculated by two-way ANOVA test with Tukey’s multiple comparisons (**c**, **f**).

### Ionizing radiation, gadolinium-based nanoparticles AGuIX and combined treatment induce AMPK activation and macrophage mitochondrial fragmentation

Considering that mitochondrial dynamics may influence the functional reprogramming of TAMs [25], we analyzed the shape of mitochondria in PMA-differentiated THP1 macrophages that were treated with single dose of 0.2 Gy IR and with indicated concentrations of AGuIX. After 1 hour of treatment, we detected the expression of the translocase of the outer mitochondrial membrane 20 (TOM20) by confocal microscopy and observed that single dose 0.2 Gy IR and treatments with 100 nM, 200 nM, 0.6 mM or 1.2 mM AGuIX led to the accumulation of shortened, rounded, fragmented mitochondria in treated PMA-differentiated THP1 macrophages (Figs. 3a-3e), as compared to control PMA-differentiated THP1 macrophages. We demonstrated that the frequency of PMA-differentiated THP1 macrophages showing mitochondrial fragmentation increased in dose-dependent manner with AGuIX concentrations (Figs. 3f and 3g). Beside, the treatments of PMA-differentiated THP1 macrophages with 0.6 mM or 1.2 mM AGuIX, respectively, triggered the accumulation of fragmented mitochondria in more than 75% to 80% of treated macrophages (Fig. 3g) as compared to control PMA-differentiated THP1 macrophages. These results thus revealed that IR and treatments with AGuIX altered mitochondrial dynamics and led to the accumulation of fragmented mitochondria in treated macrophages. Interestingly, we showed that the accumulation of fragmented mitochondria on treated PMA-differentiated THP1 macrophages was increased when macrophages were irradiated in presence of 100 nM or 200 nM AGuIX (Figs. 3a, 3b, 3e and 3f). Considering the strong effects of 0.6 mM or 1.2 mM AGuIX on the frequency of macrophages showing mitochondrial fragmentation (Fig. 3g), the combination of single dose 0.2 Gy IR with 0.6 mM or 1.2 mM AGuIX failed to show a significant enhancement of mitochondrial fragmentation in treated PMA-differentiated THP1 macrophages (Fig. 3g), as compared to irradiated PMA-differentiated THP1 macrophages. Altogether, these results demonstrate that single dose 0.2 Gy IR, treatment with different concentrations of AGuIX and combined treatments affect the mitochondrial dynamics of treated PMA-differentiated THP1 macrophages. To identify signaling pathways associated with deregulated mitochondrial dynamics, we determined whether the adenosine monophosphate (AMP) activated protein kinase (AMPK), which was recently involved in the regulation of mitochondrial dynamics [26], is phosphorylated and activated in response to these treatments. The expression of the activating phosphorylation of AMPK on threonine 172 (AMPKT172*) was then determined on control and treated PMA-differentiated THP1 macrophages with western blot analysis (Figs. 3h and 3i). Interestingly, we observed that AMPKT172* phosphorylation was strongly increased when PMA-differentiated THP1 macrophages were irradiated with all tested concentrations of AGuIX (Figs. 3h and 3i). All these processes were detected after 1 hour of treatment (Fig. 3) and before the induction of the proinflammatory activation of macrophages (Fig. 2), thus highlighting an unsuspected link between treatments with IR and/or AGuIX, the activation of AMPK, mitochondrial fragmentation and the proinflammatory reprogramming of macrophages.

**Fig. 3.**
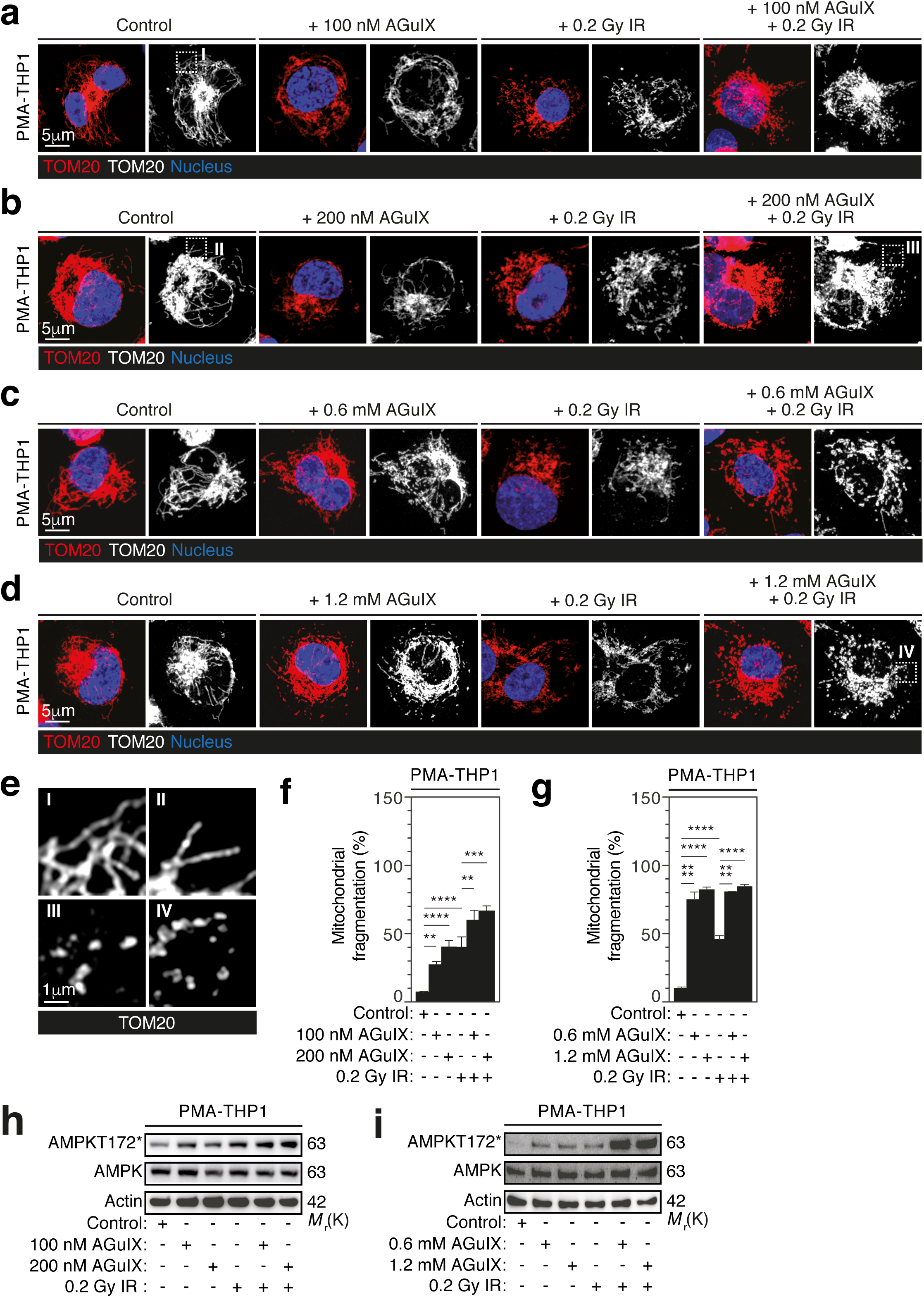
AGuIX alone and combined treatment with IR alters macrophage mitochondrial dynamics and activates AMPK. **a**-**d** Confocal micrographs of mitochondrial networks of PMA-differentiated human THP1 macrophages stained for TOM20 after 1 hour of treatment with control, 100 nM AGuIX, 0.2 Gy IR and the combination of 0.2 Gy IR with 100 nM AGuIX (**a**), with control, 200 nM AGuIX, 0.2 Gy IR and the combination of 0.2 Gy IR with 200 nM AGuIX (**b**), with control, 0.6 mM AGuIX, 0.2 Gy IR and the combination of 0.2 Gy IR with 0.6 mM AGuIX (**c**), or with control, 1.2 mM AGuIX, 0.2 Gy IR and the combination of 0.2 Gy IR with 1.2 mM AGuIX (**d**) are shown. Red and white pseudo-colors of the same confocal micrographs are shown (**a-d**). Squares indicated selected regions of interest (ROI) **I**, **II**, **III** and **IV** for magnification. **e** Higher magnifications of mitochondrial networks observed in ROI **I**, **II**, **III** and **IV** are shown. **f**, **g** Frequency of PMA-differentiated THP1 macrophages revealing mitochondrial fragmentation after 1-hour treatment with indicated conditions are shown (**f, g**). **h**, **i** AMPK and AMPK172* expressions detected on PMA-differentiated THP1 macrophages after 1-hour treatment with control, 100 nM AGuIX, 200 nM AGuIX, 0.2 Gy IR and the combination of 0.2 Gy IR with 100 nM or 200 nM AGuIX (**h**), or with control, 0.6 mM AGuIX, 1.2 mM AGuIX, 0.2 Gy IR and the combination of 0.2 Gy IR with 0.6 mM or 1.2 mM AGuIX (**i**). Actin is used as loading control. Data in **a**, **b**, **c**, **d**, **h** and **i** are representative of n=3 independent experiments. Data in **f** and **g** are means ± S.E.M from n=3 independent experiments. P-values (****P< 0.0001, ***P< 0.001 and **P< 0.01) were calculated by using one-way ANOVA test with Tukey’s multiple comparisons.

### AMPK activation controls the mitochondrial dynamics and the proinflammatory reprogramming of macrophages treated with IR, AGuIX and IR+AGuIX combination

We then investigated the effects of the AMPK activation on the proinflammatory reprogramming of macrophages induced by IR, AGuIX and combined treatment. PMA-differentiated THP1 macrophages were treated with 0.2 Gy single dose irradiation and 200 nM AGuIX in presence of 10 μM AMPK inhibitor Dorsomorphin (DRS) and analyzed for AMPKT172* expression after 1 hour treatment. As previously shown (Fig. 3h), IR and AGuIX induced AMPKT172* phosphorylation and IR+AGuIX combination strongly enhanced AMPKT172* phosphorylation (Fig. 4a). The pharmacological inhibition of AMPK with Dorsomorphin strongly impaired AMPKT172* phosphorylation (Fig. 4a) and the upregulation of IRF5 expression (Fig. 4b) that were detected respectively 1 and 48 hours after 0.2 Gy single dose irradiation of PMA-differentiated THP1 macrophages in presence of 200 nM AGuIX. These results suggest that AMPK activation plays a central role during IR-, AGuIX-, IR+AGuIX-mediated activation of anti-inflammatory macrophages toward a proinflammatory phenotype. To further characterize effects of AMPK on the proinflammatory reprogramming of macrophages in response to IR alone, AGuIX alone or AGuIX+IR combination, we then analyzed effects of the genetic depletion of AMPKα2, which is a catalytic subunit isoform of the serine/threonine kinase AMPK [27], on the mitochondrial fragmentation, the induction of ATMS1981* phosphorylation and the upregulation of IRF5 that were detected 1 hour (for mitochondrial fragmentation) and 24 hours (for ATMS1981* and IRF5) after 0.2 Gy single dose irradiation of PMA-differentiated THP1 macrophages in presence of 1.2 mM AGuIX. The specific depletion of AMPKα2 by means of small interfering RNA (Fig. 4c) significantly reduced the frequency of treated macrophages showing fragmented mitochondria (Figs. 4d and 4e), inhibited ATMS1981* phosphorylation (Fig. 4f) and impaired IRF5 upregulation (Fig. 4f). Altogether, these results confirm the essential role of AMPK during the functional reprogramming of anti-inflammatory macrophages into proinflammatory macrophages and identify signaling events (Mitochondrial fragmentation → ATMS1981* → IRF5) that are induced by AMPK activation and are required for the proinflammatory reprogramming of macrophages in response to IR, AGuIX and IR+AGuIX combination.

**Fig. 4.**
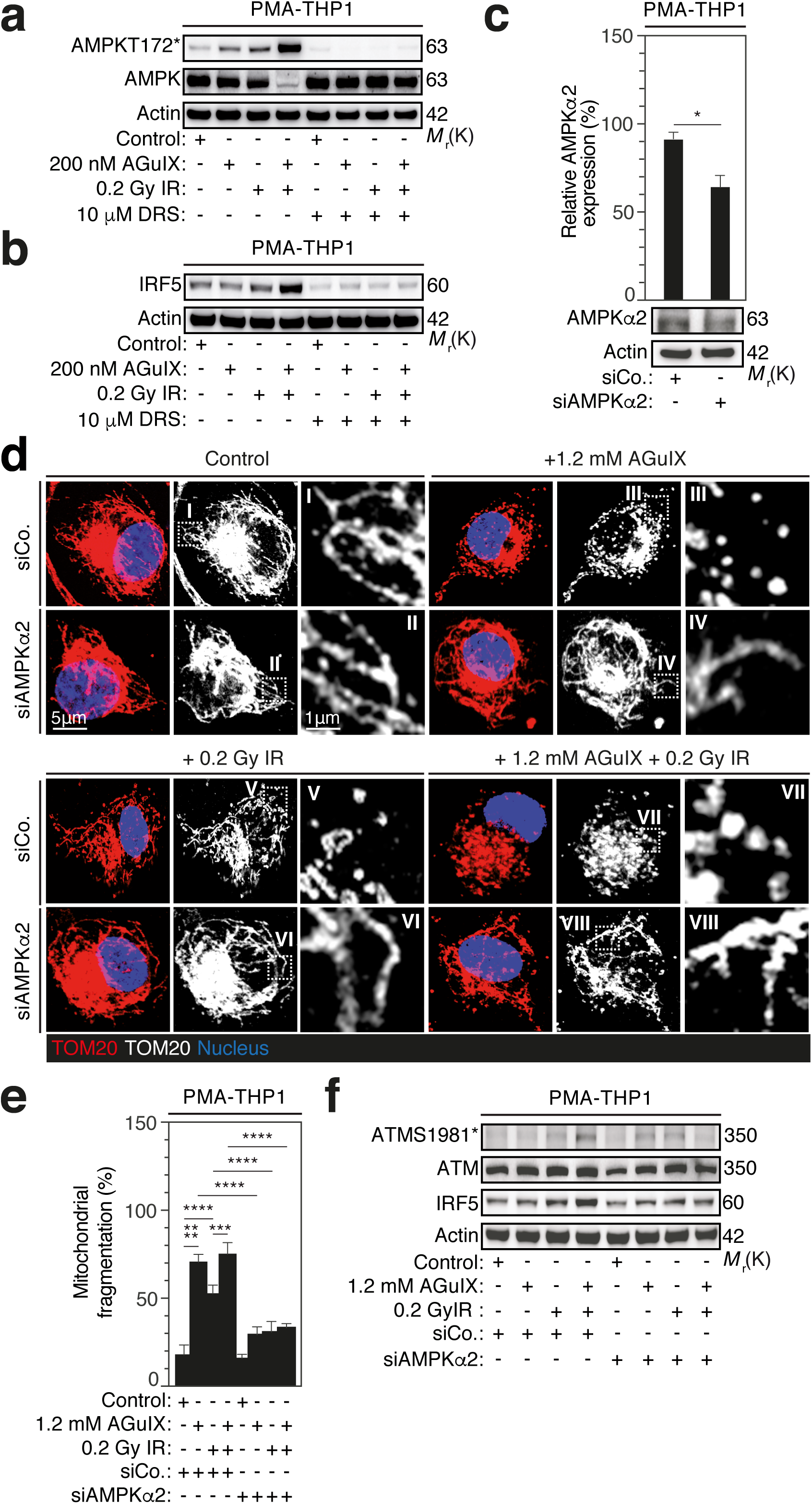
AMPK mediates mitochondrial fragmentation in response to AGuIX alone and their combination with IR. **a**, **b** AMPKT172* (**a**), AMPK (**a**) and IRF5 (**b**) expressions after 1 hour (**a**) and 48 hours (**b**) of culture of PMA-differentiated human THP1 macrophages that have been incubated with 10 µM of Dorsomorphin (DRS) and treated with control, 200 nM AGuIX, 0.2 Gy IR and the combination of 0.2 Gy IR with 100 nM or 200 nM AGuIX. Actin is used as loading control. **c** AMPKα2 expression and its quantification in PMA-differentiated human THP1 macrophages that were transfected during 48 hours with siRNAs specific for AMPKα2. Actin is used as loading control. **d**, **e** Confocal micrographs (**d**) and percentages (**e**) of PMA-THP1 macrophages showing mitochondrial fragmentation after 1-hour culture of PMA-THP1 macrophages depleted for AMPKα2 and treated with control, 1.2 mM AGuIX, 0.2 Gy IR and the combination of 0.2 Gy IR with 1.2 mM AGuIX. Red and white pseudo-colors of the same confocal microscopy images are shown (**d**). Squares indicated selected ROI **I**-**VIII** for magnification. Higher magnification details of the mitochondrial networks in ROI **I**-**VIII** are shown. **f** ATMS1981*, ATM, AMPKα2 and IRF5 expressions after 24 hours of culture of PMA-differentiated THP1 macrophages depleted for AMPKα2 and treated with control, 1.2 mM AGuIX, 0.2 Gy and the combination of 0.2 Gy IR with 1.2 mM AGuIX. Actin is used as loading control. Data in **a**, **b**, **c**, **d** and **f** are representative of n=3 independent experiments. Data in **c** and **e** are means ± S.E.M from n=3 independent experiments. P-values (****P< 0.0001, ***P< 0.001 and *P< 0.05) were calculated by using one-tailed unpaired Student’s t-test (**c**) or one-way ANOVA test with Tukey’s multiple comparisons (**e**).

### AMPK activation and proinflammatory reprogramming of TAMs are detected following IR+AGuIX combination in syngeneic mouse model of colorectal cancer

In order to investigate the ability of IR+AGuIX combination to enhance AMPK activation and thus TAM proinflammatory reprogramming *in vivo*, BALB/c mice were injected subcutaneously with CT26 cells. When the tumor volume reached 60 mm^3^, mice were intravenously injected with AGuIX (420 mg/kg) and received a single dose of local radiotherapy of 4 Gy IR (IR+AGuIX combination), were treated with control conditions (AGuIX or 4 Gy IR) or not treated. To analyze TAM reprogramming, tumor biopsies were analyzed at day 19 after treatments using confocal microscopy. The numbers of TAMs (F4/80^+^) (Figs. 5a-5c), TAMs showing AMPK activation (AMPKT172*^+^ F4/80^+^) (Figs. 5a and 5d) and TAMs expressing iNOS (iNOS^+^F4/80^+^) (Figs. 5b and 5e) detected in the tumor bed were determined. The total number of TAMs was not affected by treatments with AGuIX, IR or AGuIX+IR combination, as compared to control tumors (Figs. 5a-5c). However, a significant slight increase of TAMs number was observed in IR+AGuIX treated tumors as compared to the tumors treated with AGuIX alone (Fig. 5c). In accordance with our *in vitro* data (Fig. 3), IR induced the activating phosphorylation of AMPK (AMPKT172*) in TAMs and IR+AGuIX combination increased the number of TAMs showing AMPKT172* (AMPKT172*^+^F4/80^+^) detected in tumor beds (Figs. 5a and 5d). As we previously demonstrated [13], IR increased the number of proinflammatory activated TAMs expressing iNOS as compared to control group (Figs. 5b and 5d). More importantly, AGuIX+IR combination significantly enhanced TAM proinflammatory reprogramming, as compared to IR group (Figs. 5b and 5d), further confirming the data obtained *in vitro* (Fig. 2). Of note and in the contrary of our *in vitro* results (Figs. 2 and 3), AGuIX treatment *in vivo* did not induce AMPKT172* (Fig. 5d) and proinflammatory activation of TAMs (Fig. 5e) indicating that the combination of IR and AGuIX is required to overcome the immunosuppressive reprogramming of TAMs and to trigger their proinflammatory activation. Indeed, the accumulation of AMPKT172*^+^ TAMs detected after AGuIX+IR combination positively correlated with TAM proinflammatory reprogramming that we previously associated with the response to preoperative radiotherapy in locally advanced rectal cancer [13]. Altogether, these results demonstrate that AGuIX+IR combination may dictate the proinflammatory reprogramming of TAMs through the modulation of AMPK-dependent mitochondrial dynamic.

**Fig. 5.**
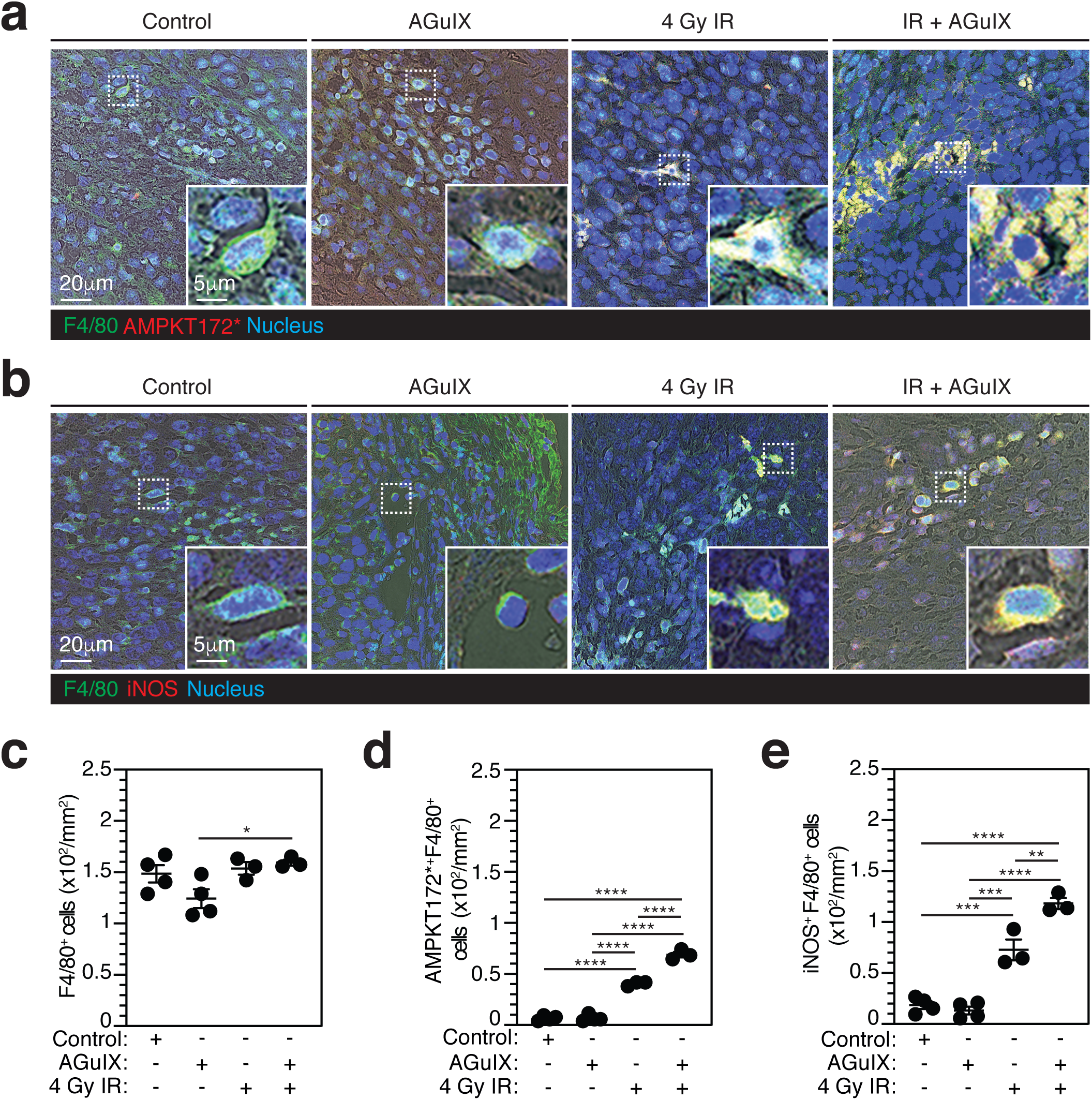
AGuIX and IR combination triggers AMPK activation and proinflammatory reprogramming in TAMs. **a-e** Confocal micrographs (**a**, **b**) and counts (**c**-**e**) of F4/80^+^ (**a**-**c**), AMPKT172*^+^F4/80^+^ (**a**, **d**) and iNOS^+^F4/80^+^ (**b**, **e**) tumor-associated macrophages detected on tumor biopsies obtained from CT26 tumor bearing BALB/c mice after 19 days of treatment with control, AGuIX (420 mg/kg), 4 Gy IR and IR+AGuIX combination. Squares indicated ROI for magnification. Inserts show magnifications of selected ROI. Contrast phase revealed tumor cellularity. Data shown in **a** and **b** are representative for n=4 mice (for control and AGuIX) and n=3 mice (for IR and AGuIX+IR). Data in **c**, **d** and **e** are means ± S.E.M from n=4 mice (for control and AGuIX) and n=3 mice (for IR and AGuIX+IR). P-values (****P< 0.0001, ***P<0.001, **P<0.01 and *P<0.05) were calculated by using one-way ANOVA test with Tukey’s multiple comparisons.

## Discussion

Resistance to cancer treatments is a major obstacle to overcome for cancer cure. The therapeutic targeting of tumor microenvironment (TME) has recently emerged as a promising approach to reduce refractoriness to cancer treatments and to enhance cancer treatment efficacy [1]. TAMs, which are major cellular components of TME, have gained attention as novel cellular targets for anticancer therapy. In the vast majority of cancers, TAMs are associated with poor prognosis and therapeutic resistances [4, 28, 29]. Several therapeutic approaches targeting TAMs for depletion, repression of migration and/or functional reprogramming have been approved or are still under consideration for clinical use [29]. These strategies include neutralizing antibodies or small molecules inhibitors of colony-stimulating factor 1 receptor (CSF1R) [30, 31], CC-motif chemokine ligand 2 (CCL2) [32], CC-chemokine receptor 2 (CCR2) [33], surface receptor TREM2 [34] and PI3Kγ [35]. We previously demonstrated that radiotherapy also triggers the proinflammatory reprogramming of TAMs through the induction of a molecular cascade that requires the increased expression of NOX2, the production of ROS, the promotion of double strand breaks, the activation and the phosphorylation ATMS1981*, the induction of IRF5, the expression of iNOS and the secretion of proinflammatory cytokines IL-1β, IL-6, IL-8, IFN-γ-α [13]. In addition, we revealed that other cancer treatments such as cisplatin and the poly(ADP-ribose)polymerase (PARP) inhibitor olaparib also induced the proinflammatory activation of macrophages by stimulating this signaling pathway [13], thus revealing that combining radiotherapy with other modalities of cancer treatments might enhance the efficacy of radiotherapy through the reprogramming of TAMs.

In this study, we demonstrated that the combination of radiotherapy with gadolinium-based nanoparticles AGuIX enhances the ability of macrophages to undergo proinflammatory activation in response to IR (Fig. 6). Consistent with several *in vitro* and *in vivo* studies showing that IR+AGuIX combination increases the dose deposition in several cancer cells [36, 37], our results revealed that the *in vitro* treatment of macrophages with IR+AGuIX combination is associated with an increased dose deposition in treated macrophages, as compared to irradiated macrophages, thus providing the first evidence that IR+AGuIX combination is able to target and potentially control the fate of TAMs. Moreover, our results showed that the kinase ATM is phosphorylated on serine 1981 and activated in response to IR+AGuIX combination (Fig. 1), thus suggesting that the kinase ATM may potentially play a central role during IR+AGuIX-mediated proinflammatory macrophage reprogramming, as we previously shown for IR alone [13]. Our results revealed that the transcription factor IRF5, which is the major transcriptional regulator of IR-mediated proinflammatory macrophage activation [13], exhibited an increased expression in response to IR+AGuIX combination. The increased secretion of IL-1β and IL-6, and an increased production of iNOS were also detected in response to IR+AGuIX combination, as compared to treatments with IR or AGuIX alone, demonstrating that IR+AGuIX combination enhanced the ability of macrophages to be reprogrammed into proinflammatory macrophages. These results are consistent with previous publications revealing the functional switch of macrophages from anti-inflammatory to proinflammatory phenotype in response to other metallic nanoparticles alone (such as gold-based nanoparticles [38], iron oxide nanoparticles [39] and titanium dioxide nanoparticles [40]) or combined treatments with IR [41].

**Fig. 6.**
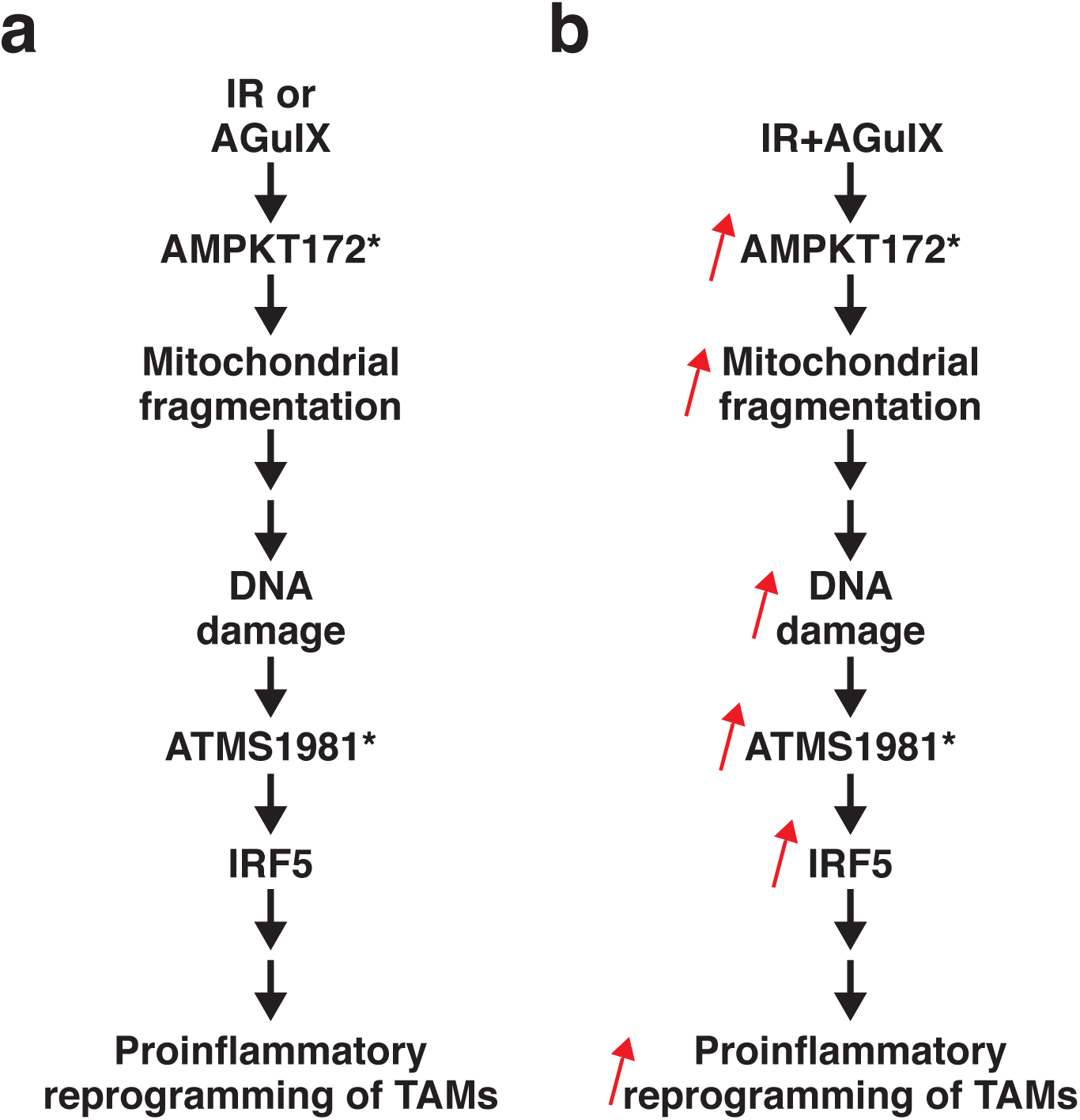
Proposed model for the roles of AMPK and mitochondrial fragmentation in the proinflammatory macrophage reprogramming induced by AGuIX and AGuIX+ IR combination.

In this study, we also demonstrated that all treatments used (IR, AGuIX and IR+AGuIX combination) induced the phosphorylation and the activation of the AMPK, which is a central metabolic sensor that regulates cellular energy homeostasis through the modulation of numerous cellular processes such as autophagy [42], oxidative phosphorylation [43], mitochondrial dynamics and integrity [44]. We showed that AMPKT172* is rapidly detected in response to treatments and positively correlated with the accumulation of fragmented “stressed” mitochondria in the cytosol of treated macrophages. Interestingly, we demonstrated that pharmacological inhibition (with DRS) or genetic depletion of AMPKα2 abolished mitochondrial fragmentation and impaired the proinflammatory reprogramming of treated macrophages, thus revealing a counterintuitive link between the AMPK activation, the mitochondrial fragmentation and the proinflammatory reprogramming of macrophages. In one hand, the activation of AMPK was previously involved in the promotion of the anti-inflammatory reprogramming of macrophages [45] and the repression of LPS-induced proinflammatory reprogramming of astrocytes, microglia and peritoneal macrophages [46]. The activation of AMPK was also shown to increase fatty acid oxidation [47]. In other hand, the activation of AMPK is required for the fragmentation of mitochondria through the modulation of the dynamin-related protein 1 (DRP1) activity [44] and the mitochondrial fragmentation was associated metabolic regulation of several immune cells such as lymphocytes, regulatory T cells [48] and NK cells [49].

Our results provide the first evidence that the activation of AMPK by IR modulates mitochondrial dynamics and activates ATM for the proinflammatory reprogramming of macrophages. Further molecular characterizations are needed to precise the contribution of AMPKT172* and mitochondrial fragmentation to the proinflammatory signaling pathway elicited by IR that we previously described [13].

Importantly, we confirmed using a murine CT26 colorectal carcinoma preclinical model that IR+AGuIX combination enhanced the activation of AMPK and the proinflammatory reprogramming of TAMs, as compared to what we observed on biopsies obtained from tumor-bearing mice receiving radiotherapy alone. In contrary to what we reported *in vitro*, AGuIX alone did not trigger the activation of AMPK and promote the proinflammatory activation of TAMs. These results revealed that in absence of radiotherapy, AGuIX is not sufficient to overcome the anti-inflammatory activation of TAMs and highlighted the rational to combine AGuIX with radiotherapy *in vivo* to sensitize anti-inflammatory TAMs to radiotherapy and promote their proinflammatory activation.

### Conclusions

One of the major therapeutic approaches to boost antitumor immune response is to overcome the immunosuppressive TME in which anti-inflammatory TAMs are the pillars. Here we demonstrate that the combination of IR with AGuIX enhanced the ability of anti-inflammatory macrophages to undergo proinflammatory activation in response to IR and might be useful to improve patient’s response to radiotherapy through an AMPK-dependent proinflammatory reprogramming of TAMs.

## Materials and methods

### Cells and Reagents

The human THP1 monocytes (#TIB-202, ATCC, USA) and murine RAW264.7 macrophages (#TIB-71, ATCC, USA) were maintained in RMPI-1640-Glutamax medium (#61870044) and DMEM-GlutaMax (#31966047), respectively, both from Life Technologies (Carlsbad, CA, USA), supplemented with 10% heat-inactivated fetal bovine serum (FBS) (Eurobio, #CVFSVF0001, France) and 100 IU/ml penicillin–streptomycin (Life Technologies, #15140130, USA). THP1 macrophages were obtained after differentiation of THP1 monocytes with 320 nM PMA (#tlrl-PMA, Invivogen, San Diego, CA, USA) during 24 hours. After PMA removal, treated cells were rested for additional 24 hours. All cells were maintained under 5% CO2 humidified atmosphere at 37 °C. Dorsomorphin (#3093/10) was from Tocris Bioscience (Bristol, UK). PMA (#tlrl-PMA) was from Invivogen. Gadolinium (Gd)-based AGuIX nanoparticles were synthesized and provided by NH TherAguix SA company (Lyon, France). AGuIX nanoparticles were suspended at 100 mg/ml in RNAse/DNAse free water before used.

### Irradiation and treatment with gadolinium-nanoparticle AGuIX

THP1 macrophages were differentiated in 12-well plates (10^6^ cells per well), 24-well plates (5x10^5^ cells per well) or 48-well plates (2.5x10^5^ cells per well) as described above. RAW264.7 macrophages were seeded in 24-well plates (5x10^4^ per well) 24 hours before treatments. Cells were then incubated with different concentrations of AGuIX diluted in HBSS (#14025092, Thermo Fisher Scientific, USA) at 37 °C for 1 hour and irradiated immediately or after one hour incubation in medium at the indicated doses with X-ray irradiator (1 Gy/min, X-RAD 320, Precision X-Ray, Madison, USA). Nanoparticles suspension was removed and macrophages were incubated in medium supplemented with 10% FBS during 30 min, 1 hour, 24 hours and 48 hours after irradiation for western blot analysis or for immunofluorescence microscopy. Cell culture supernatants used for western blot analysis were collected 48 hours after treatments.

### RNA-mediated interference

The SMARTpool siGENOME Human PRKAA2 (5563) siRNAs (M-005361-02-0005) against AMPK alpha-2 and siGENOME non-targeting siRNA Pool #1 (D-001206-13-05) as control were purchased from Dharmacon (Lafayette, CO, USA). The sequences of SMARTpool siGENOME Human PRKAA2 (5563) siRNA are: siRNA-1: 5’- GUACCUACGUUAUUUAAGA-3’; siRNA-2: 5’-GGAAGGUAGUGAAUGCAUA-3’; siRNA-3: 5’-GACAGAAGAUUCGCAGUUU-3’; siRNA-4: 5’- ACAGAAGAUUCGCAGUUUA-3’. A pool of four on-target plus non-targeting siRNAs from siGENOME was used as control siRNA. PMA-differentiated THP1 macrophages were transfected using a double transfection with Lipofectamine RNAi max (#13778150, Thermo Fisher Scientific, USA) according to the manufacturer’s instructions. Briefly, THP1 monocytes were differentiated as described above (10^6^ cells/ml/well in 12-well plate). Lipofectamine RNAi max and siRNA were mixed in Opti-MEM reduced serum medium (#31985-070, Thermo Fisher Scientific, USA) and incubated at room temperature for 5 minutes. Then, transfection mix was added to THP1 macrophages with a final siRNA concentration of 12 nM and cells were incubated at 37°C for 24 hours. Macrophages were then re-transfected with siRNA with the same concentration as used for the first transfection and incubated at 37°C for additional 24 hours.

### Antibodies

Antibodies used for immunofluorescence microscopy were anti-phospho-H2AX (Ser139*) (#05-636, diluted at 1:100) antibody from EMD Millipore (Billerica, MA, USA), anti-AMPK alpha (Phospho Thr172) (AMPKT172*) (#GTX52341, diluted at 1:50) and anti-phospho-ATM (Ser1981*) (#GTX132146, diluted at 1:100) antibodies from Genetex (Irvine, USA), anti-iNOS (#ab3523, diluted at 1:500 for immunofluorescence labelling of fixed cells and 1:20 for tumor biopsies) and anti-F4/80 (#ab6640, diluted at 1:50) antibodies from Abcam (Cambridge, UK), and anti-Tom20 (F-10) (#sc-17764, diluted at 1:100) antibody from Santa Cruz Biotechnology (Oregon, USA). Antibodies used for western blots were anti-phospho-ATM (Ser1981) (10H1.E12) (#4526, diluted at 1:1000), anti-ATM (D2E2) (#2873, diluted at 1:1000), anti-phospho-AMPKα (Thr172) (40H9) (#2535, diluted at 1:1000), anti-AMPKα (#2532, diluted at 1:1000) antibodies from Cell Signaling Technology (Massachusetts, USA), anti-IRF5 (#ab21689, diluted at 1:40000), anti-beta Actin HRP [AC-15] (#ab49900, diluted at 1:10000), anti-IL-1 beta (#ab2105, diluted at 1:1000) antibodies from Abcam (Cambridge, UK), anti-AMPKα2 (#AF2850, diluted at 1:1000) and anti-IL6 (#AB-206-NA, diluted at 1:500) antibodies from R&D Systems (Minneapolis, MN, USA). Secondary antibodies used for immunofluorescence labelling of fixed cells were goat anti-rabbit antibodies conjugated to Alexa Fluor 488 (#11034, diluted at 1:500) or Alexa Fluor 546 (#A11010, diluted at 1:500) and goat anti-mouse antibodies conjugated to Alexa Fluor 488 (#11001, diluted at 1:500) or Alexa Fluor 546 (#11030, diluted at 1:500) from Thermo Fisher Scientific (USA). For immunofluorescence labelling of tumor biopsies, the secondary antibodies used were Alexa Fluor 488 goat anti-rat (#A11006, diluted at 1:500) and Alexa Fluor 546 goat anti-rabbit (#A11035, diluted at 1:500) antibodies from Thermo Fisher Scientific (USA). The secondary antibodies used for western blot analysis were goat anti-rabbit conjugated to horseradish peroxidase-conjugated (HRP) (#4050-05, diluted at 1:1000) and anti-mouse-HRP (#1031-05, diluted at 1:1000) from SouthernBiotech (USA).

### Immunofluorescence microscopy

THP1 macrophages were treated and incubated as described above on 10 mm coverslips (#41001110, Deckglaser, Germany) at a concentration of 2.5x10^5^ cells per well in 48-well plates. RAW264.7 macrophages were seeded 24 hours before treatments on 13 mm coverslips (#ECN631-1578, VWR, France) in 24-well plates (5x10^4^ per well). Macrophages were fixed with 4% (w/v) paraformaldehyde (#1.00496.5000, Sigma-Aldrich, Germany) for 10 minutes, permeabilized phosphate buffered saline (PBS) containing 0.3% Triton X-100 (Sigma-Aldrich, Germany) for 20 minutes, blocked with PBS containing 10% FBS (10% FBS-PBS) for 1 hour. Primary antibodies in 10% FBS-PBS were incubated for 1 hour and 30 minutes at room temperature. Primary antibodies were used at dilution of 1/100 excepted for anti-iNOS antibody (1:500) and anti-ATMS1981* (1:50). Then, macrophages were incubated with secondary antibodies conjugated with Alexa Fluor-488 green or Alexa Fluor-546 red (Thermo Fisher Scientific, USA) (1:500) and Hoechst 33342 (#H3570, Thermo Fisher Scientific, USA) (dilution 1:1000) in 10% FBS-PBS during 30 min at room temperature. Coverslips were mounted with Fluoromount-G mounting medium (#00-4958-02, Thermo Fisher Scientific, USA) on microscope cover glasses (#631-0170, VWR, France). The quantifications (with a minimum of 300 cells for each condition) were performed with Leica DMI8 fluorescent microscopy (Leica Microsystems, Nanterre, France) using a 63X oil objective. For representative images shown in figures, images were acquired by confocal microscopy (SP8, Leica Microsystems, Nanterre, France) by hybrid detectors (pinhole airy: 0.6; pixel size: 180 nm) with optimal optical sectioning (OOS) of 0.8 µm from the top to the bottom of each cell. Images were generated on max-intensity of z projection images using Image J software (NIH, USA). The staining of mouse tumor sections (4 µm/section) was performed as we previously described [13, 50, 51]. Briefly, after deparaffinization and antigen retrieval (in 10 mM sodium citrate buffer, pH 6 at 96 °C for 30 min), mouse tumor sections were blocked for 1 hour in 10% FBS-PBS, and incubated overnight at 4°C with the anti-F4/80 (#ab6640, 1:50) and/or anti-iNOS (#ab3523, 1:20) antibodies from Abcam (Cambridge, UK), and/or anti-AMPK alpha (Phospho Thr172) (AMPKT172*) antibody (#GTX52341, 1:50) from Genetex (Irvine, USA) in 10% FBS-PBS. Then, mouse tumor sections were incubated for 2 hours at room temperature with Alexa Fluor 488 goat anti-rat (#A11006, 1:500) antibodies, Alexa Fluor 546 goat anti-rabbit (#A11035, 1:500) antibodies and Hoechst 33342 (#H3570, 1:1000) from Thermo Fisher Scientific (USA). Mouse tumor sections were mounted on microscope cover glasses (#631-0170, VWR, France) using Fluoromount-G (#00-4958-02, Thermo Fisher Scientific, USA). Total number of F4/80^+^ TAMs expressing iNOS or AMPKT172* was determined on at least nine fields (209.5 µm x 209.5 µm) acquired with SP8 (Leica) confocal microscope and was normalized to tumor tissue surface.

### Western blot analysis

Total cellular proteins were extracted in CHAPS buffer containing 10 mM TRIS (pH = 7.4), 850 mM NaCl, and 0.1% CHAPS hydrate (C3023, Sigma-Aldrich, Germany) and complemented with protease (#4693124001) and phosphatase (#04906837001) inhibitors from Roche (Basel, Switzerland) during 20 minutes. About 3-30 μg of protein extract were run on 4-12% (#NP0332BOX), 10% (#NP0302BOX) or 12% (#NP0342BOX) NuPAGE gels (Invitrogen, Illkrich, France) and then, were transferred onto a 0.2 µm nitrocellulose membrane (#162-0112, Bio-Rad, Marnes-la-coquette, France) at room temperature. After blocking with 5% bovine serum albumin (BSA) in Tris-buffered saline and 0.1% Tween 20 at room temperature for 1 hour, membranes were incubated with primary antibodies overnight at 4 °C and during 2 hours at room temperature with horseradish peroxidase (HRP)-conjugated anti-mouse or anti-rabbit IgG (SouthernBiotech, USA). Membranes were then revealed using G:BOX Chemi XL1.4 Fluorescent & Chemiluminescent Imaging System (Syngene, Cambridge, UK).

### *In vivo* studies

Mouse experiments were performed in compliance with French and European laws and regulations. The local institutional animal ethics board and French ethical committee (CEEA26) of the Ministère de l’éducation nationale, de l’enseignement supérieur et de la recherche approved mouse experiments (permission number: 2020-064-27337). Experiments were performed in accordance with the international and national recommendations for proper use and care for the laboratory animals. BALB/c female mice at the age of 6-7 weeks (purchased from Janvier Laboratories (France) and maintained in the animal facility of Institut Gustave Roussy (France)) were injected subcutaneously with 5x10^5^ CT26 cells suspended in 50 μl cold PBS. When tumor volumes reached approximately 60 mm^3^, mice were randomized based on their tumor volumes and body weights in four groups. Mice were then either intravenously injected with AGuIX nanoparticles (420 mg/Kg), irradiated with the single dose of 4 Gy IR at the tumor bed using Varian Tube NDI 226 (USA) or Gulmay X-ray irradiator (250 kV, tube current of 15 mM, beam filter of 0.2 mm Cu) (Gulmay Medical L.T.D, UK) with a dose rate of 1.08 Gy/min, treated with AGuIX and IR combination or only injected with AGuIX vehicle (RNAse/DNAse free water). At nineteen days, mice were anesthetized using isoflurane gas. Tumors were excised, fixed in 10 % formalin solution (#1.00496.5000, Sigma-Aldrich, Germany), embedded in paraffin and cut into 4 µm thick sections for immunofluorescence microscopy analysis.

### Statistical analysis

Statistical analyses were performed with GraphPad Prism version 6.0b (GraphPad Software, La Jolla, CA, USA) using two-way ANOVA test with Tukey’s multiple comparisons (for Figs. 1c, 1f, 2c and 2f), one-way ANOVA test with Tukey’s multiple comparisons (for Figs. 1g, 1i, 3f, 3g, 4e, 5c, 5d and 5e) or one-tailed unpaired Student’s t-test (for Fig. 4c). The data represented in the figures are means ± SEM from independent experiments. The statistical significances illustrated on the figure graphs are indicated as *P<0.05, **P<0.01, ***P<0.001, and ****P<0.0001. The n size, the statistical tests and the multiple comparisons adjustments are indicated in the corresponding figure legends.

## Acknowledgements

This work has benefited from the facilities and expertise of the Imaging and Cytometry Platform UMS 3655 CNRS / US 23 INSERM, Gustave Roussy Cancer Campus Villejuif, France. Z.M. and C.M. are recipients of PhD fellowships from Université Paris-Saclay. D.T. is recipient of PhD fellowship from ANRT (Cifre#2019/0639 with NH TherAguix SA). A.M.-K. is recipient of PhD fellowship from Ecole et Loisir. T.T.C.-P. is supported by a grant from Institut National du Cancer (INCa 16087).

## Author Contributions

J.-L.P. designed the study. Z.M., D.T., A.M.-K., T.T.C.-P. and A.A. performed experiments. Z.M., A.A. and J.-L.P analyzed the results and wrote the paper. Z.M. and A.A. assembled the figures and performed statistical analysis. A.M.-K., T.T.C.-P., C.L., S.B., A.P., S.D., T.D, F.L., O.T. and G.L.D. provided advices and edited the paper.

## Availability of data and materials

The data needed to evaluate the conclusions of the paper are present in the paper.

## Declarations

### Ethics approval and consent to participate

Mouse experiments were performed in compliance with French and European laws and regulations. The local institutional animal ethics board and French ethical committee (CEEA26) of the Ministère de l’éducation nationale, de l’enseignement supérieur et de la recherche approved mouse experiments (permission number: 2020-064-27337). Experiments were performed in accordance with the international and national recommendations for proper use and care for the laboratory animals.

## References

1. Bejarano, L., M.J.C. Jordao, and J.A. Joyce, Therapeutic Targeting of the Tumor Microenvironment. Cancer Discov, 2021. 11(4): p. 933–959.

2. Kloosterman, D.J. and L. Akkari, Macrophages at the interface of the co-evolving cancer ecosystem. Cell, 2023. 186(8): p. 1627–1651.

3. Guerriero, J.L., Macrophages: The Road Less Traveled, Changing Anticancer Therapy. Trends Mol Med, 2018. 24(5): p. 472–489.

4. Mantovani, A., et al., Tumour-associated macrophages as treatment targets in oncology. Nat Rev Clin Oncol, 2017. 14(7): p. 399–416.

5. Qiu, Y., et al., Next frontier in tumor immunotherapy: macrophage-mediated immune evasion. Biomark Res, 2021. 9(1): p. 72.

6. Cassetta, L. and J.W. Pollard, A timeline of tumour-associated macrophage biology. Nat Rev Cancer, 2023. 23(4): p. 238–257.

7. Biswas, S.K. and A. Mantovani, Macrophage plasticity and interaction with lymphocyte subsets: cancer as a paradigm. Nat Immunol, 2010. 11(10): p. 889–96.

8. Pathria, P., T.L. Louis, and J.A. Varner, Targeting Tumor-Associated Macrophages in Cancer. Trends Immunol, 2019. 40(4): p. 310–327.

9. Feng, M., et al., Phagocytosis checkpoints as new targets for cancer immunotherapy. Nat Rev Cancer, 2019. 19(10): p. 568–586.

10. Bruni, D., H.K. Angell, and J. Galon, The immune contexture and Immunoscore in cancer prognosis and therapeutic efficacy. Nat Rev Cancer, 2020. 20(11): p. 662–680.

11. Noy, R. and J.W. Pollard, Tumor-associated macrophages: from mechanisms to therapy. Immunity, 2014. 41(1): p. 49–61.

12. Wu, Q., et al., Modulating Both Tumor Cell Death and Innate Immunity Is Essential for Improving Radiation Therapy Effectiveness. Front Immunol, 2017. 8: p. 613.

13. Wu, Q., et al., NOX2-dependent ATM kinase activation dictates pro-inflammatory macrophage phenotype and improves effectiveness to radiation therapy. Cell Death Differ, 2017. 24(9): p. 1632–1644.

14. Hainfeld, J.F., et al., Gold nanoparticles: a new X-ray contrast agent. Br J Radiol, 2006. 79(939): p. 248–53.

15. Dorsey, J.F., et al., Gold nanoparticles in radiation research: potential applications for imaging and radiosensitization. Transl Cancer Res, 2013. 2(4): p. 280–291.

16. Taupin, F., et al., Gadolinium nanoparticles and contrast agent as radiation sensitizers. Phys Med Biol, 2015. 60(11): p. 4449–64.

17. McQuaid, H.N., et al., Imaging and radiation effects of gold nanoparticles in tumour cells. Sci Rep, 2016. 6: p. 19442.

18. Wang, H., et al., Cancer Radiosensitizers. Trends Pharmacol Sci, 2018. 39(1): p. 24–48.

19. Le Duc, G., et al., Toward an image-guided microbeam radiation therapy using gadolinium-based nanoparticles. ACS Nano, 2011. 5(12): p. 9566–74.

20. Dufort, S., et al., Nebulized gadolinium-based nanoparticles: a theranostic approach for lung tumor imaging and radiosensitization. Small, 2015. 11(2): p. 215–21.

21. Lux, F., et al., AGuIX((R)) from bench to bedside-Transfer of an ultrasmall theranostic gadolinium-based nanoparticle to clinical medicine. Br J Radiol, 2019. 92(1093): p. 20180365.

22. Lakshmanan, V.K., et al., Nanomedicine-based cancer immunotherapy: recent trends and future perspectives. Cancer Gene Ther, 2021.

23. Irvine, D.J. and E.L. Dane, Enhancing cancer immunotherapy with nanomedicine. Nat Rev Immunol, 2020. 20(5): p. 321–334.

24. Du, Y., et al., Radiosensitization Effect of AGuIX, a Gadolinium-Based Nanoparticle, in Nonsmall Cell Lung Cancer. ACS Appl Mater Interfaces, 2020. 12(51): p. 56874–56885.

25. Gao, Z., et al., Mitochondrial dynamics controls anti-tumour innate immunity by regulating CHIP-IRF1 axis stability. Nat Commun, 2017. 8(1): p. 1805.

26. Herzig, S. and R.J. Shaw, AMPK: guardian of metabolism and mitochondrial homeostasis. Nat Rev Mol Cell Biol, 2018. 19(2): p. 121–135.

27. Hawley, S.A., et al., Characterization of the AMP-activated protein kinase kinase from rat liver and identification of threonine 172 as the major site at which it phosphorylates AMP-activated protein kinase. J Biol Chem, 1996. 271(44): p. 27879–87.

28. DeNardo, D.G. and B. Ruffell, Macrophages as regulators of tumour immunity and immunotherapy. Nat Rev Immunol, 2019. 19(6): p. 369–382.

29. Cassetta, L. and J.W. Pollard, Targeting macrophages: therapeutic approaches in cancer. Nat Rev Drug Discov, 2018. 17(12): p. 887–904.

30. Tap, W.D., et al., Structure-Guided Blockade of CSF1R Kinase in Tenosynovial Giant-Cell Tumor. N Engl J Med, 2015. 373(5): p. 428–37.

31. Moughon, D.L., et al., Macrophage Blockade Using CSF1R Inhibitors Reverses the Vascular Leakage Underlying Malignant Ascites in Late-Stage Epithelial Ovarian Cancer. Cancer Res, 2015. 75(22): p. 4742–52.

32. Loberg, R.D., et al., Targeting CCL2 with systemic delivery of neutralizing antibodies induces prostate cancer tumor regression in vivo. Cancer Res, 2007. 67(19): p. 9417–24.

33. Yang, Z., et al., CCL2/CCR2 Axis Promotes the Progression of Salivary Adenoid Cystic Carcinoma via Recruiting and Reprogramming the Tumor-Associated Macrophages. Front Oncol, 2019. 9: p. 231.

34. Molgora, M., et al., TREM2 Modulation Remodels the Tumor Myeloid Landscape Enhancing Anti-PD-1 Immunotherapy. Cell, 2020. 182(4): p. 886–900 e17.

35. Kaneda, M.M., et al., Macrophage PI3Kgamma Drives Pancreatic Ductal Adenocarcinoma Progression. Cancer Discov, 2016. 6(8): p. 870–85.

36. Hu, P., et al., Gadolinium-Based Nanoparticles for Theranostic MRI-Guided Radiosensitization in Hepatocellular Carcinoma. Front Bioeng Biotechnol, 2019. 7: p. 368.

37. Sancey, L., et al., The use of theranostic gadolinium-based nanoprobes to improve radiotherapy efficacy. Br J Radiol, 2014. 87(1041): p. 20140134.

38. Yen, H.J., S.H. Hsu, and C.L. Tsai, Cytotoxicity and immunological response of gold and silver nanoparticles of different sizes. Small, 2009. 5(13): p. 1553–61.

39. Zanganeh, S., et al., Iron oxide nanoparticles inhibit tumour growth by inducing pro-inflammatory macrophage polarization in tumour tissues. Nat Nanotechnol, 2016. 11(11): p. 986–994.

40. Zhao, F., et al., Titanium dioxide nanoparticle stimulating pro-inflammatory responses in vitro and in vivo for inhibited cancer metastasis. Life Sci, 2018. 202: p. 44–51.

41. Jin, J.Z., Q., Engineering nanoparticles to reprogram radiotherapy and immunotherapy: recent advances and future challenges J Nanobiotechnology, 2020. 18(1): p. 75.

42. Kim, J., et al., AMPK and mTOR regulate autophagy through direct phosphorylation of Ulk1. Nat Cell Biol, 2011. 13(2): p. 132–41.

43. Tailor, D., et al., Novel Aza-podophyllotoxin derivative induces oxidative phosphorylation and cell death via AMPK activation in triple-negative breast cancer. Br J Cancer, 2021. 124(3): p. 604–615.

44. Toyama, E.Q., et al., *Metabolism.* AMP-activated protein kinase mediates mitochondrial fission in response to energy stress. Science, 2016. 351(6270): p. 275–281.

45. Sag, D., et al., Adenosine 5’-monophosphate-activated protein kinase promotes macrophage polarization to an anti-inflammatory functional phenotype. J Immunol, 2008. 181(12): p. 8633–41.

46. Giri, S., et al., 5-aminoimidazole-4-carboxamide-1-beta-4-ribofuranoside inhibits proinflammatory response in glial cells: a possible role of AMP-activated protein kinase. J Neurosci, 2004. 24(2): p. 479–87.

47. Pilon, G., P. Dallaire, and A. Marette, Inhibition of inducible nitric-oxide synthase by activators of AMP-activated protein kinase: a new mechanism of action of insulin-sensitizing drugs. J Biol Chem, 2004. 279(20): p. 20767–74.

48. Field, C.S., et al., Mitochondrial Integrity Regulated by Lipid Metabolism Is a Cell-Intrinsic Checkpoint for Treg Suppressive Function. Cell Metab, 2020. 31(2): p. 422–437 e5.

49. Zheng, X., et al., Mitochondrial fragmentation limits NK cell-based tumor immunosurveillance. Nat Immunol, 2019. 20(12): p. 1656–1667.

50. Allouch, A., et al., SUGT1 controls susceptibility to HIV-1 infection by stabilizing microtubule plus-ends. Cell Death Differ, 2020. 27(12): p. 3243–3257.

51. Allouch, A., et al., CDKN1A is a target for phagocytosis-mediated cellular immunotherapy in acute leukemia. Nat Commun, 2022. 13(1): p. 6739.

